# The SUMO protease Ulp2 regulates genome stability and drug resistance in the human fungal pathogen *Candida albicans*

**DOI:** 10.1101/2021.12.06.471441

**Authors:** Marzia Rizzo, Natthapon Soisangwan, Jan Soetaert, Samuel Vega-Estevez, Anna Selmecki, Alessia Buscaino

## Abstract

Stress-induced genome instability in microbial organisms is emerging as a critical regulatory mechanism for driving rapid and reversible adaption to drastic environmental changes. In *Candida albicans*, a human fungal pathogen that causes life-threatening infections, genome plasticity confers increased virulence and antifungal drug resistance. Discovering the mechanisms regulating *C. albicans* genome plasticity is a priority to understand how this and other microbial pathogens establish life-threatening infections and develop resistance to antifungal drugs. We identified the SUMO protease Ulp2 as a critical regulator of *C. albicans* genome integrity through genetic screening. Deletion of *ULP2* leads to hypersensitivity to genotoxic agents and increased genome instability. This increased genome diversity causes reduced fitness under standard laboratory growth conditions but enhances adaptation to stress, making *ulp2Δ/Δ* cells more likely to thrive in the presence of antifungal drugs. Whole-genome sequencing indicates that *ulp2Δ/Δ* cells counteract antifungal drug-induced stress by developing segmental aneuploidies of chromosome R and chromosome I. We demonstrate that intrachromosomal repetitive elements drive the formation of complex novel genotypes with adaptive power.

## Introduction

Understanding how organisms survive and thrive in changing environments is a fundamental question in biology. Genetic variation is central to environmental adaptation as it allows selection of certain genotypes better fit to grow in new environments. Different types of genetic change contribute to genetic variability, including *(i)* mutations such as single-base alteration and small (<100 bp) insertions or deletions (indels), *(ii)* large (>1 kb) deletions and duplications, *(iii)* whole-chromosome or segmental-chromosome aneuploidy and *(iv)* translocations and complex genomic rearrangements [1]. Furthermore, diploid cells can undergo Loss of Heterozygosity (LOH) driven by cross-overs or gene conversions between the two homologous chromosomes [2]. Excessive genome instability is harmful in the absence of selective pressure as it alters the copy-number of many genes, leading to unbalanced protein levels [3]. However, an unstable genome can provide rapid adaptive power in hostile environments [4,5] because it provides genetic diversity upon which selection can act.

Genome plasticity – the ability to generate genomic variation – is emerging as a critical adaptive mechanism in human microbial pathogens that need to adapt rapidly to extreme environmental shifts, including changes in temperature, pH and nutrient availability following colonisation of different host environments [6,7]. One such organism is *Candida albicans*, the most common human fungal pathogen and the most prevalent cause of death due to fungal infection. *C. albicans* is part of the normal microflora of most healthy individuals where it colonises the skin, mucosal surface, gastrointestinal and the female genitourinary tract. However, *C. albicans* can become a dangerous pathogen causing a wide range of infections, from superficial mucosal infections to life-threatening disseminated diseases [8]. Azole antifungal agents, such as Fluconazole (FLC), are the most commonly prescribed drugs for treating *Candida* infections [9–11]. FLC targets the enzyme lanosterol 14α-demethylase, encoded by *ERG11*, blocking biosynthesis of ergosterol, an essential component of the fungal cell membrane [12,13]. As a result, FLC arrests *C. albicans* cell growth without killing the fungus. This fungistatic, rather than fungicidal, mode of action allows for the evolution of drug-resistant strains [14]. One primary mechanism of drug resistance is an increased production of the FLC target, Erg11 enzyme, diluting the activity of the drug [12]. This high target production is often due to increased activity of the transcription factor Upc2 activating *ERG11* transcription [15– 18]. Overproduction of efflux pumps, such as the *C. albicans* proteins Cdr1, Cdr2 and Mdr1, can also drive FLC resistance by decreasing intracellular FLC levels [19]. In recent years, genome plasticity has emerged as a critical adaptive mechanism causing antifungal drug resistance. *C. albicans* is a diploid organism with a highly heterozygous genome organised into 2 × 8 chromosomes (2n = 16) [20,21]. Population studies have identified a remarkable genomic variation among *C. albicans* isolates and specific chromosomal variations are selected during host-niche colonisation [22–28]. Indeed, many drug-resistant isolates exhibit karyotypic diversity, including aneuploidy and gross chromosomal rearrangements that can confer resistance due increased copy number of specific genes including *ERG11*, and/or multidrug transporters [7,29,30].

*C. albicans* genome instability is not random: it occurs more frequently at specific hotspots that are often repetitive [31–35]. Subtelomeric regions and the *rDNA* locus are among the most unstable genomic sites [34,36]. Indeed, *C. albicans* subtelomeric regions are enriched in transposons-derived repetitive sequences and protein-coding genes [31,37]. Most notable are the <u>t</u>e<u>lo</u>mere-associated *TLO* genes, a family of 14 closely related paralogues encoding proteins similar to the Mediator 2 subunit of the mediator transcriptional regulator [38–40]. The majority of *TLO* genes are located at subtelomeric regions except *TLO34*, located at an internal locus on the left arm of Chr1 [38]. The number and position of *TLO* genes vary widely between clinical isolates, indicating significant plasticity with potential consequences for the fitness of the organism [34]. The *rDNA* locus consists of a tandem array of a ∼12□kb unit repeated 50 to 200 times on chromosome R; rDNA length polymorphisms occur frequently [21,34]. In addition to these complex repetitive elements, different types of Long Repeat Sequences (65 bp to 6.5 Kb) dispersed across the *C. albicans* genomes have been shown to drive karyotype variation during adaptation to antifungal drugs and passage through the mouse host [32,33]. *C. albicans* genome plasticity is regulated by environmental conditions: the genome is relatively stable under optimal laboratory growth conditions but becomes more unstable under stress conditions [41,42]. For example, FLC treatment drives a global increase in LOH, chromosome rearrangements and aneuploidy [41,42]. This increased genetic variation facilitates selection of fitter genotypes [28,29]. Similarly, higher rates of genomic variation are detected following passage of *C. albicans in vivo* relative to passage *in vitro* [35,43]. It is unknown if and how stress regulates genome plasticity. The discovery of such regulatory mechanisms will be essential to reveal how resistance to antifungal drugs emerges.

This study posits that gene deletions for critical regulators of *C. albicans* genome integrity would cause higher genome variation and rapid adaptation to FLC. To test this hypothesis, we performed a genetic screen to identify modulators of *C. albicans* genome stability. The screen led to the identification of the SUMO protease Ulp2. In the absence of stress, *ULP2* deletion leads to elevated genome instability causing fitness defects and hypersensitivity to genotoxic agents. In contrast, the elevated genome instability of the *ulp2 Δ/Δ* strain is advantageous in the presence of high FLC doses. This is because the increased genetic diversity expands the pool of genotypes upon which selection can act, driving adaptation to a new stress environment (FLC), concomitantly rescuing the fitness defects associated with *ULP2* deletion. We also demonstrate that intrachromosomal repetitive elements are sites of genetic diversity that drive the formation of complex novel genotypes with adaptive potential.

## Results

### A systematic genetic screen identifies the Ulp2 as a regulator of C. albicans genotoxic stress response

To identify factors regulating *C. albicans* genome integrity, we utilised a deletion library comprising a subset (674/3000) of *C. albicans* genes that are not conserved in other organisms or have a functional motif potentially related to virulence [44]. As defects in genome integrity lead to hypersensitivity to genotoxic agents [45], the deletion library was screened for hypersensitivity to two DNA damaging agents: Ultraviolet (UV) irradiation which induces formation of pyrimidine dimers [46], and Methyl MethaneSulfonate (MMS), which leads to replication blocks and base mispairing [47].

Genotoxic stress hypersensitivity was semi-quantitatively scored by comparing the growth of treated versus untreated on a scale of 0 to 4, where 0 indicates no sensitivity, and 4 specifies strong hypersensitivity (**Fig 1A)**. The screen identified 32 gene deletions linked to DNA damage hypersensitivity (UV or MMS score ≥2). Almost half of these hits (14/32; ∼44%) are genes predicted to encode components of the DNA Damage Response pathway (7/32; ∼22%) or the cell division machinery (7/32; ∼22%) (**Table S1**). For example, the top 4 hits of the screen were *MEC3, RAD18, GRR1* and *KIP3* genes. Although *C. albicans MEC3* and *RAD18* are uncharacterised, they encode for proteins, conserved in other organisms, that are universally involved in sensing DNA damage (Mec3) [48] and in DNA post-replication repair (Rad18) [49]. *C. albicans GRR1* and *KIP3* are required for cell cycle progression [50] and mitotic spindle organisation, respectively [51] (**Fig 1B** and **Table S1**). ∼25% (8/32) of the remaining hits are genes encoding proteins with no apparent orthologous in the two well-studied yeast model systems (*S. cerevisiae* and *S. pombe*). This high percentage is not surprising as one of the criteria used to select target genes for the deletion library was the lack of conservation between *C. albicans* and yeast model systems [44]. The remaining 10 hits are genes encoding for proteins with diverse functions, including stress response (*HOG1*) [52], transcriptional and chromatin regulation (*SPT8*, SET3) [53–55], transport (*YPT7, DUR35, NPR2, FCY2*) [56–59], protein folding (*HCH1*) [60], MAP kinase pathway (*STT4*) [61] and cell wall biosynthesis (*KRE5*) [62].

**Fig 1.**
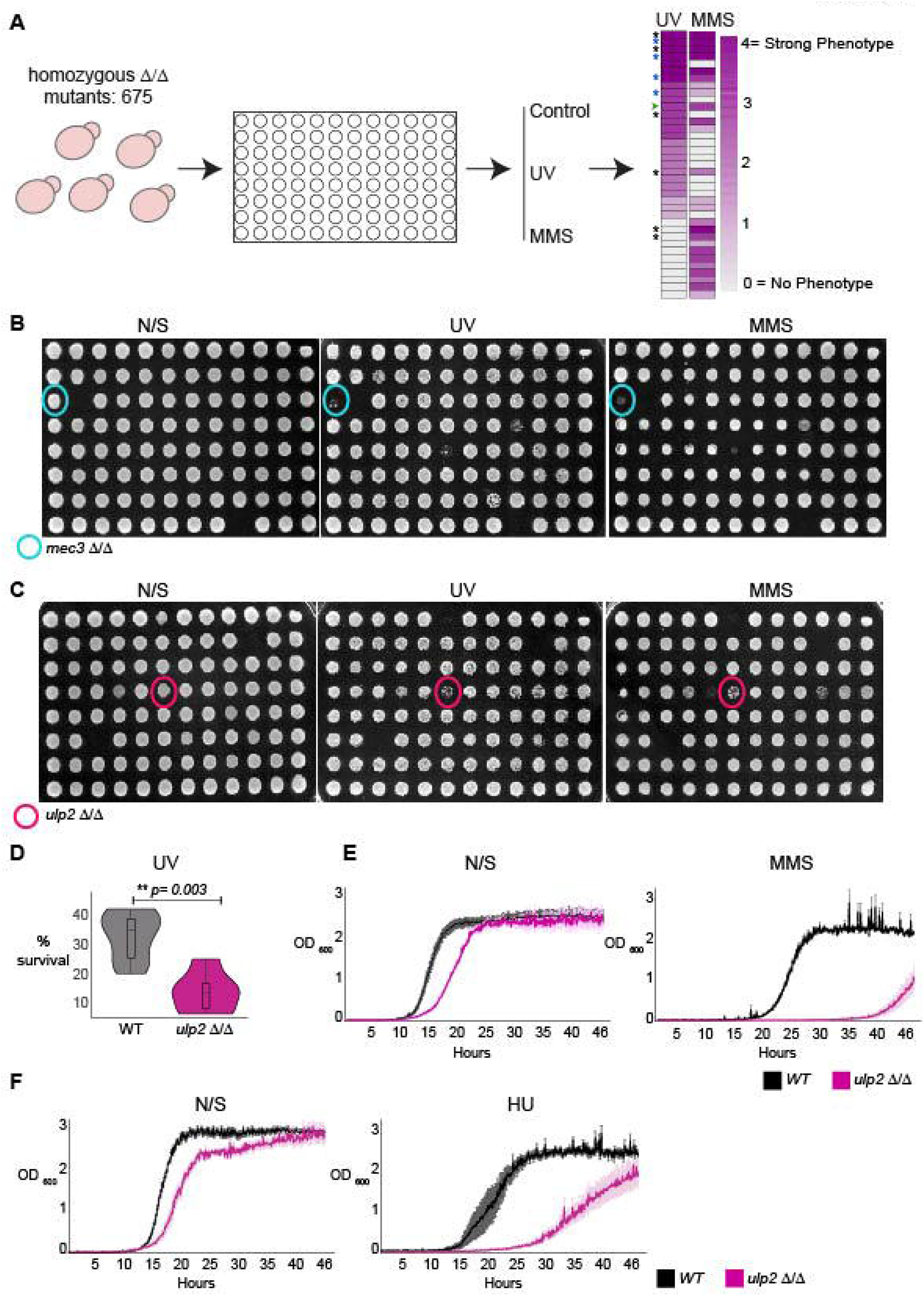
ULP2 is a regulator of C. albicans genotoxic stress response. **(A)** Schematic representation of the screening strategy. 674 *C. albicans* deletion strains were screened using a 96-plate format for hypersensitivity to UV and MMS. Hypersensitivity was scored by comparing the growth of treated vs untreated on a scale of 0 (white) to 4 (magenta). Black *: genes encoding for DNA damage and sensing repair pathway components, Blue *: genes encoding for cell division and chromosome segregation machinery, Green arrow: *ulp2 Δ/Δ* **(B)** Data for a plate containing *mec3 Δ/Δ* strain (cyan circle). Growth on Non-selective (N/S) media or following UV and MMS treatment is shown. **(C)** Data for a plate containing *ulp2 Δ/Δ* strain (magenta circle). Growth on Non-selective (N/S) media or following UV and MMS treatment is shown **(D)** Colony-forming Unit assay of UV treated WT and *ulp2 Δ/Δ* strains. % survival is shown. **(E)** Growth curve on WT and *ulp2 Δ/Δ* strains grown in non-selective (N/S) liquid media and MMS-containing liquid media. Error bars: standard deviation (SD) of three biological replicates **(F)** Growth curve on WT and *ulp2 Δ/Δ* strains grown in non-selective (N/S) liquid media and HU-containing liquid media.

One of the highest-ranked genes on our screen is *ULP2* (CR_03820C/ *orf19*.*4353:* EMS score:3, UV score:3) encoding for a SUMO protease (**Fig 1C**). SUMOylation is a dynamic and reversible post-translation modification in which a member of the SUMO family of proteins is conjugated to target proteins at lysine residues by E1 activating enzymes, E2 conjugating enzymes and E3 ligases [63–65]. SUMO proteases remove the polypeptide SUMO from target proteins, regulating their function, activity or localisation [66,67].

*C. albicans ULP2* is an excellent candidate for a modulator of stress-induced genome plasticity for several reasons: *(i)* <u>p</u>ost-<u>t</u>ranslation <u>m</u>odifications (PTMs), such as SUMOyolation, are rapid and reversible. Consequently, PTMs can modulate genome instability in response to rapid and transient environmental changes [68,69], *(ii)* protein sumoylation is emerging as a critical stress response mechanism across eukaryotes [66,70–73] *(iii) C. albicans* protein sumoylation levels change in response to environmental stresses encountered in the host [74].

Colony-Forming Unit (CFU) assays of UV-treated cells confirm the importance of *ULP2* in DNA damage resistance as UV treatment reduced the number of CFU in *a ulp2 Δ/Δ* strain (∼14.5% survival) compared to a wild-type (WT) strain (∼33.7% survival:) (**Fig 1D**). Furthermore, the *ulp2* Δ/Δ strain also displayed a reduced growth rate in liquid media containing MMS or Hydroxyurea (HU), a chemotherapeutic agent that challenges genome integrity by stalling replication forks [75] (**Fig 1E** and **1F**). Thus, *ULP2* has a role in the response to a wide range of genotoxic agents.

### ULP2 but not ULP1 is required for survival under stress

*C. albicans* contains three putative SUMO-deconjugating enzymes: Ulp1, Ulp2 and Ulp3 (**Fig 2A**). Sequence comparison between the three *C. albicans* Ulp proteins and the two *S. cerevisiae* Ulps (Ulp1 and Ulp2) reveals that although the *C. albicans* proteins are poorly conserved, the amino acid residues essential for catalytic activity are conserved. This analysis suggests that all *C. albicans* Ulps are active SUMO proteases (**Fig 2A** and **2B**). Accordingly, recombinantly expressed *C. albicans* Ulp1, Ulp2 and Ulp3 have SUMO-processing activity *in vitro* [76]. Similarly to *S. cerevisiae ULP1, C. albicans ULP3* is an essential gene and was not investigated further in this study [77].

**Fig 2.**
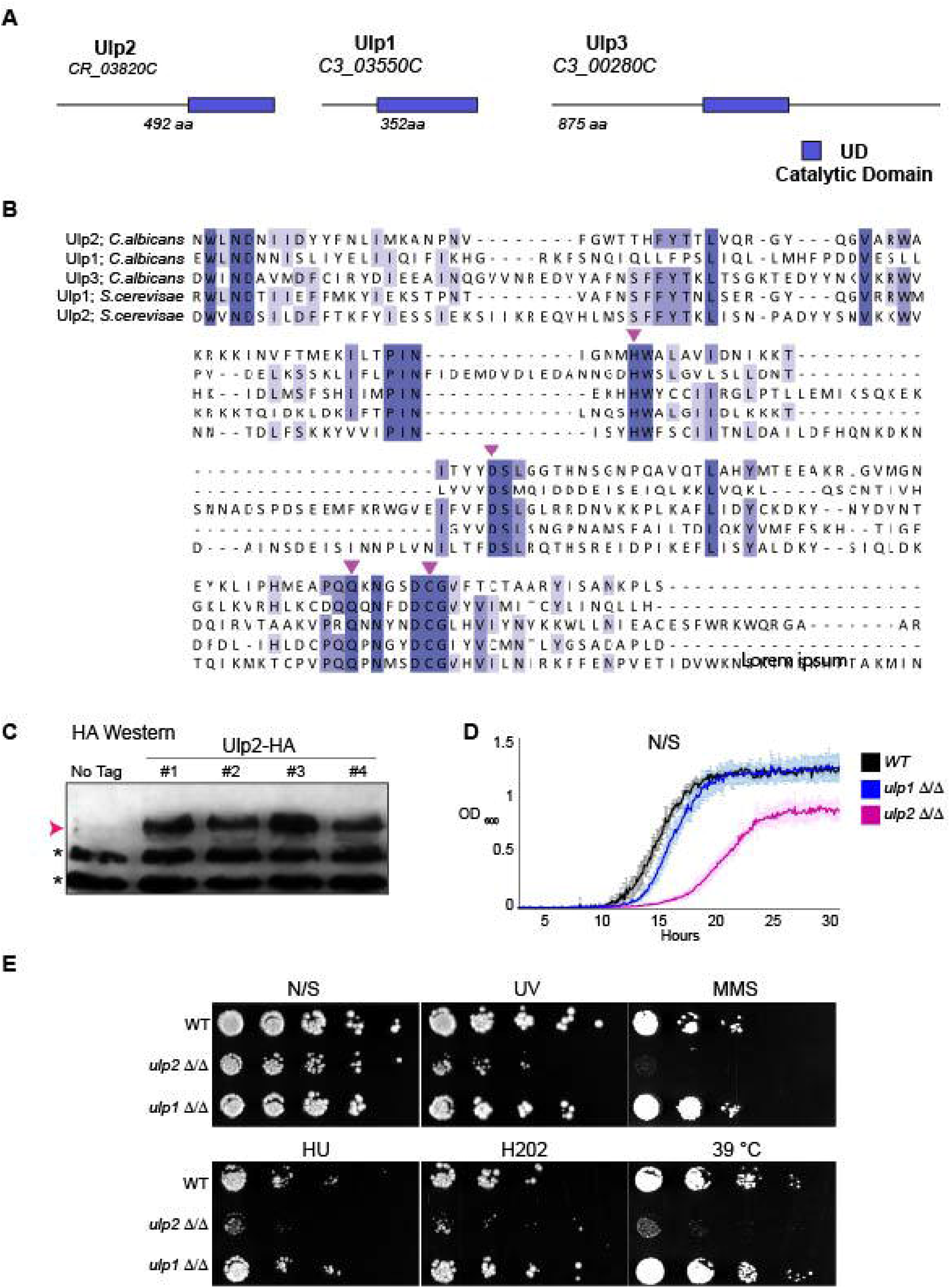
ULP2 is necessary for survival under stress. **(A)** Schematic representation of Ulp1, Ulp2 and Ulp3 protein organisation. The systematic name and the amino acid (aa) number is indicated for each protein. Blue box: putative catalytic UD domain typical of Ulp SUMO proteases **(B)** Protein alignments of the three *C. albicans* Ulp proteins (Ulp1, Ulp2 and Ulp3) and the two *S. cerevisiae* proteins (Ulp1 and Ulp2). Magenta arrows: amino acids essential for SUMO protease activity **(C)** HA Western Blot analysis of 4 independent ULP2-HA integrants and the progenitor untagged control (No Tag). Magenta arrow: Ulp2-HA (Magenta arrow). *: non-specific cross-reacting bands serving as a loading control **(C)** Growth curves of WT, *ulp1 Δ/Δ* and *ulp2 Δ/Δ* strains grown in non-selective (N/S) liquid media. Error bars: standard deviation (SD) of three biological replicates **(D)** Serial dilution assay of WT, *ulp1 Δ/Δ* and *ulp2 Δ/Δ* strains grown in unstressed (N/S) or stress (UV, MMS, HU, H202 and 39 °C) growth conditions.

Previous studies suggested that *C. albicans* Ulp2 is an unstable or a very low abundant protein undetectable by Western blot analysis [76]. We reassessed *ULP2* expression by generating strains expressing, at the endogenous locus, an epitope-tagged Ulp2 protein (Ulp2-HA). Western analyses show that Ulp2-HA expression is readily detected in extracts from independent integrant strains. (**Fig 2C**). Thus, a stable Ulp2 protein is expressed in cells grown under standard laboratory growth conditions (YPD, 30 °C). To assess whether *C. albicans ULP1* and *ULP2* gene share a similar function, we engineered homozygous deletion strains for *ULP1* (*ulp1*Δ/Δ) and *ULP2* (*ulp2*Δ/Δ). Growth analysis demonstrated that deletion of *ULP2* reduces fitness as the newly generated *ulp2*Δ/Δ strain is viable, but cells are slow-growing (**Fig 2D** and **2E**). In contrast, the *ulp1*Δ/Δ strain grows similarly to the WT control in solid and liquid media (**Fig 2D** and **2E**). Phenotypic analysis confirms that *ULP2* is an important regulator of *C. albicans* stress response as, similarly to the deletion library mutant, the newly generated *ulp2* Δ/Δ strain is sensitive to different stress conditions including treatment with DNA damaging agents (UV and MMS), DNA replication inhibitor (HU), oxidative stress (H_2_0_2_) and high temperature (39°C) (**Fig 2E**) In contrast, deletion of *ULP1* did not cause any sensitivity to the tested stress conditions (**Fig 2E**).

In summary, we could not detect any phenotype associated with deletion of *ULP1*, while loss of *ULP2* leads to poor growth in standard laboratory growth conditions and hypersensitivity to multiple stresses.

### Loss of ULP2 leads to increased genome instability

To assess whether the hypersensitivity to DNA damage agents observed in the *ulp2*Δ/Δ strain was indeed due to enhanced genome instability, we deleted the *ULP2* gene from a set of tester strains containing a heterozygous *URA3*^*+*^ marker gene inserted in three different chromosomes (Chr 1, 3 and 7) [41]. We quantified the frequency of *URA3*^*+*^ marker loss by plating on plates containing the *URA3* counter-selective drug FOA and scoring the number of colonies able to grow on FOA-containing media compared to non-selective (N/S) media. Deletion of *ULP2* leads to a dramatic increase in LOH rate at all three chromosomes (Chr1: ∼5000X, Chr3: ∼18X, Chr7: ∼170X), indicating that *ULP2* is required for maintaining genome stability across the *C. albicans* genome (**Fig 3A**).

**Fig 3.**
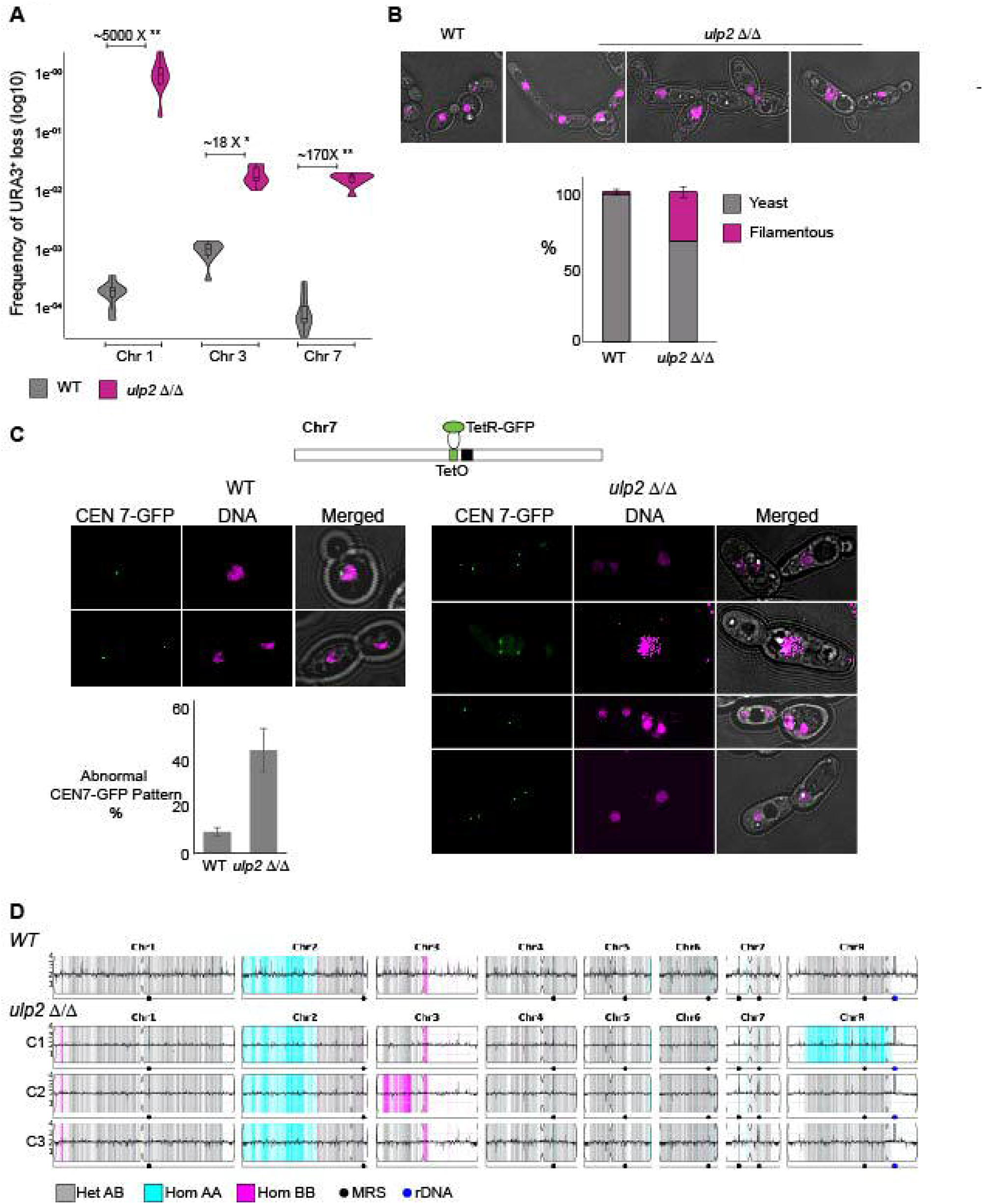
Loss of ULP2 leads to increased genome instability. **(A)** Quantification of loss of a heterozygous *URA3*^*+*^ marker gene inserted in Chr1, Chr3 and Chr7 in WT and *ulp2 Δ/Δ* strain. A fold difference of *URA3*^*+*^ marker loss between *ulp2 Δ/Δ* and WT strains is indicated. **: Chr1 (4.11E-07) and Chr7 (6.74E-05) p-value, **\*:** Chr3 (2.87E-02) p-value **(B)** *Top:* Representative images displaying the morphologies of WT and *ulp2 Δ/Δ* strains. *Bottom:* Quantification (%) of yeast and filamentous (hyphae + pseudohyphae) cells in WT and *ulp2 Δ/Δ* strains. Error bar: Standard deviation of 3 biological replicates. **(C)** *Top:* schematics of the CEN7 TetO and TetR-GFP system. *Bottom*: nuclear morphology and segregation pattern of centromere 7 (*CEN7*) in WT and *ulp2 Δ/Δ* strain. Quantification (%) of abnormal GFP-CEN7 patterns is indicated. Error bar: Standard deviation of 3 biological replicates. **(D)** Whole genome sequencing analysis of the progenitor (SN152) and three single colonies C1, C2, and C3. Data were plotted as the log2 ratio and converted to chromosome copy number (y-axis, 1-4 copies) as a function of chromosome position (x-axis, Chr1-ChrR) using the Yeast Mapping Analysis Pipeline (YMAP) [112]. Heterozygous (AB) regions are indicated with grey shading, and homozygous regions (loss of heterozygosity) are indicated by shading of the remaining haplotype, either AA (cyan) or BB (magenta). Two homozygous positions are present in the progenitor (the left side of Chr2 and a small region near the centromere of Chr3), while C1 and C2 underwent loss of heterozygosity of ChrR and Chr3.

In *C. albicans*, hypersensitivity to genotoxic stress often correlates with filamentous growth [45,78–81]. Accordingly, and in agreement with a significant role for *ULP2* in genotoxic stress response, the *ulp2*Δ/Δ strain displays a higher frequency of abnormal morphologies than a WT strain, including filamentous pseudohyphal-like and hyphal-like cells (**Fig 3B**). To assess whether the exacerbated *ulp2Δ/Δ* genome instability is linked to defective chromosome segregation, we deleted the *ULP2* gene in a reporter strain in which *TetO* sequences are integrated adjacent to the centromere (*CEN7*) of one Chromosome 7 homolog and TetR-GFP fusion protein is expressed from an intergenic region [82]. Binding of TetR-GFP to *tetO* sequences allowed visualisation of Chr7 duplication and segregation during the cell cycle. We found that deletion of *ULP2* leads to abnormal Chr7 segregation, including cells with no TetR-GFP signals or multiple TetR-GFP-foci, which was ∼5 fold higher in the *ulp2 Δ/Δ* strain than the WT control strain (**Fig 3C**).

Previous studies performed in the model system *S. cerevisiae* demonstrated that loss of *ULP2* leads to the accumulation of a specific multichromosome aneuploidy (amplification of both ChrI and ChrXII) that rescues the potential lethal defects of *ulp2* deletion by amplification of specific genes on both chromosomes [83,84]. To determine whether loss of *C. albicans ULP2* results in a specific aneuploidy, we sequenced the genome of 3 randomly selected *ulp2* Δ/Δ colonies by whole genome sequencing (WGS) and compared their genome sequences to the *C. albicans* reference genome. This analysis demonstrates that deletion of *C. albicans ULP2* does not select for specific chromosome rearrangements and identifies different genomic variations that are not present in the parental WT strain (**Fig 3D** and **Table S2**) [85]. While deletion of *ULP2* leads to very few (<10) *de novo* mutations (**Table S2**), two of the three colonies underwent extensive LOH on different chromosomes (**Fig 3D and Table S2**). For example, chromosome missegregation followed by reduplication of the remaining homologue is detected on isolate C1 (C1: ChrR) and the genome of C2 contains a long-track LOH (C2:Chr 3L) that occurred within 4.6 kb of a repeat locus on Chr3L (*PGA18*, [32]) (**Fig 3D**). Our analysis collectively demonstrates that deletion of *C. albicans ULP2* leads to increased genome instability via the formation of extensive chromosomal variation.

### Loss of ULP2 leads to drug resistance via selection of novel genotypes

We hypothesised that the increased genome instability of the *ulp2 Δ/Δ* strain would facilitate adaptation to hostile environments via selection of fitter genotypes. To test this hypothesis, we assessed whether WT and *ulp2Δ/Δ* strains differ in their ability to overcome the stress imposed by low or high concentrations of 2 drugs: Fluconazole (FLC) and caffeine (CAF). FLC was chosen because it is the most used antifungal drug in the clinic. CAF was chosen because it is associated with well-known resistance mechanisms [86,87]. Serial dilution analyses demonstrate that the *ulp2Δ/Δ* strain is not sensitive to a low FLC (15 μg/ml) dose while it is sensitive a low CAFF (5mM) doses (**Fig 4A** and **4B**).

**Fig 4.**
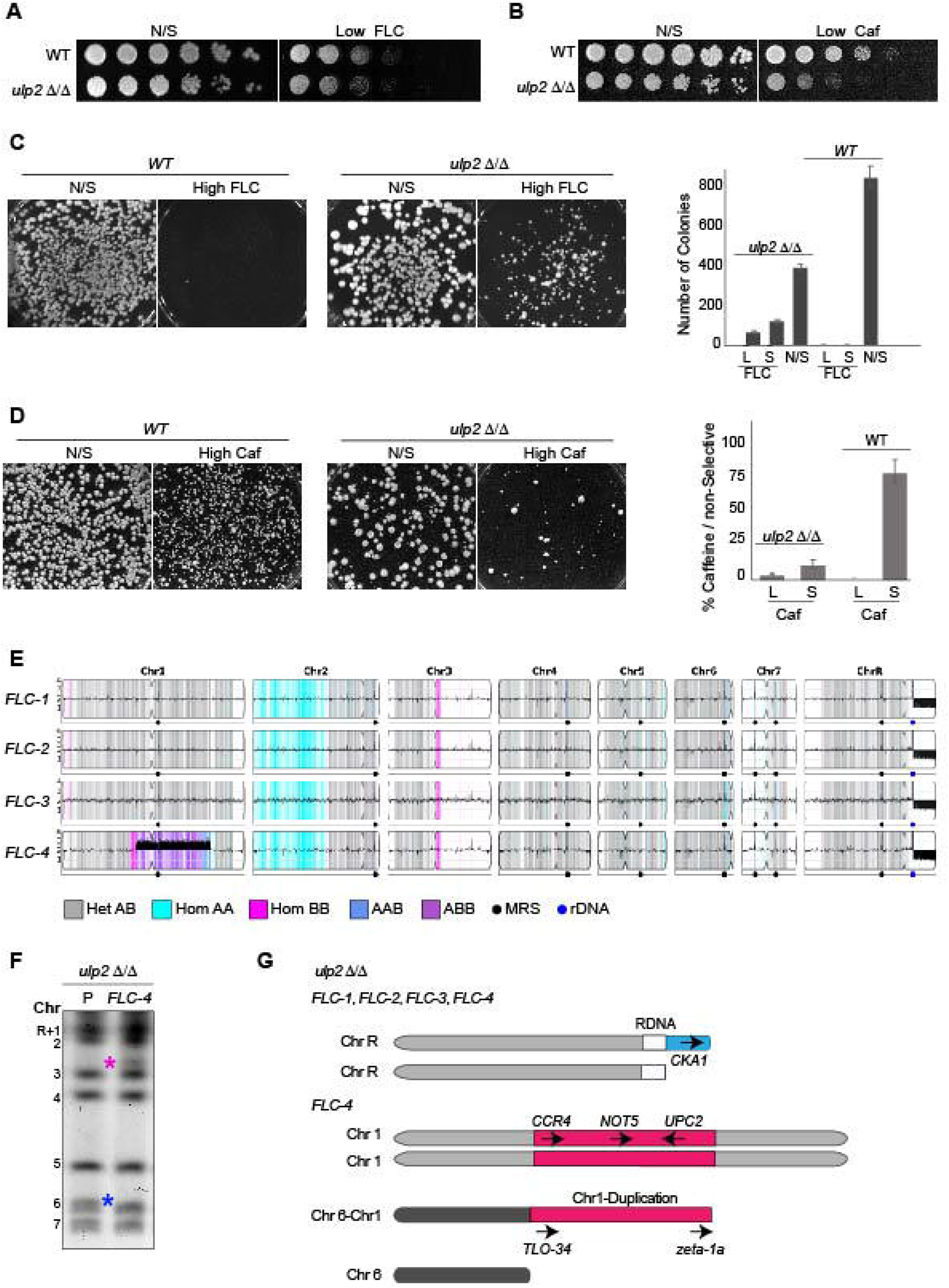
Loss of ULP2 leads to drug resistance via selection of novel genotypes. **(A)** Serial dilution assay of WT and *ulp2 Δ/Δ* strains grown in non-selective (N/S) or media containing low (15 μg/ml) concentration of fluconazole (FLC). **(B)** Serial dilution assay of WT and *ulp2 Δ/Δ* strains grown in non-selective (N/S) or media containing low (5 mM) Caffeine (CAF). **(C)** *Left:* Plating assay of *ulp2 Δ/Δ* and WT strain in media containing high (128 μg/ml) concentration of fluconazole (FLC) or non-selective (NS) media *Right:* Plating assay quantification. The number of large (L) and small (S) colonies recovered from fluconazole (FLC) containing media and non-selective (N/S) media is shown for WT and *ulp2 Δ/Δ* strains. **(D)** *Left:* Plating assay of *ulp2 Δ/Δ* and WT strain in media containing high (12 mM) concentration of caffeine (CAF) and non-selective (NS) media *Right:* Plating assay quantification. The number of large (L) and small (S) colonies recovered from caffeine (CAF)-containing media and non-selective (N/S) media is shown for WT and *ulp2 Δ/Δ strains*. **(E)** Whole genome sequencing data plotted as in Figure 3D for four single colonies isolated from 128 μg/ml fluconazole plates (*FLC1*-*FLC4*). The chromosome copy number is plotted along the y-axis (1-4 copies). All four single colonies have a recurrent segmental deletion of part of ChrRR. Colony *FLC-4* has an amplification of the middle part of Chr1. Copy number breakpoints and allele ratio changes in *FLC-4* are indicated in Figure S3. **(F)** CHEF karyotype gel stained with ethidium bromide of *ulp2 Δ/Δ* progenitor and *FLC-4* isolate. A band (blue *) corresponding to Chr6 is present in the *ulp2 Δ/Δ* progenitor and absent in the *FLC-4* isolate. Conversely, a new band (magenta *) is present in the *FLC-4* isolate but absent in the *ulp2 Δ/Δ* progenitor. **(G)** Schematics of segmental aneuploidies detected in *FLC-1, FLC-2, FLC-3* and *FLC-4* isolates.

In contrast, deletion of *ULP2* increases adaptation to high doses FLC and CAF. On plates containing an inhibitory concentration of FLC (128 μg/ml), a WT strain produced only tiny abortive colonies while the *ulp2Δ/Δ* strain produces colonies of heterogenous size (Large and Small, **Fig 4C**). The starting *ulp2Δ/Δ* strain is highly sensitive to 12 mM CAF (**Fig S1A**), and therefore a reduced number of *ulp2Δ/Δ* colonies grew at this high drug concentration compared to the WT strain (**Fig 4D**). Despite this difference, the *ulp2 Δ/Δ* strain, but not the WT strain, produces large colonies that can grow on high CAF concentration following passaging in the absence of the drug, indicative of adaptation (**Fig 4D, Fig S1B**). Thus, deletion of *ULP2* accelerates adaptation to lethal drug concentration.

To test whether enhanced drug adaption was linked with selection of novel genotypes, we sequenced the genome of 4 independent *ulp2Δ/Δ* FLC-adapted isolates (*FLC-1, FLC-2, FLC-3* and *FLC-4*). *FLC-1, FLC-2* and *FLC-3* were randomly selected from the High FLC plates and sequenced immediately. In contrast, *FLC-4* was selected because this isolate was still able to grow on high FLC following passaging in non-selective (N/S) media (**Fig S1C**). To assess for genotype heterogeneity, three *FLC-4* derived single colonies (*FLC-4a, b* and *c*) were sequenced (**Fig S2A** and **B**). The WGS analysis demonstrates that all FLC-adapted colonies have a genotype that is distinct from the *ulp2 Δ/Δ* progenitor. We detected very few (<10) *de novo* point mutations, and none of these are common among all the sequenced FLC isolates (**Table S3**). In contrast, all colonies are marked by an extensive segmental chromosome aneuploidy: a partial deletion (∼ 388 Kb) of the right arm of Chromosome R (ChrRR-Deletion). ChrRR-deletion occurs at the ribosomal DNA (25S subunit) and it extends to the right telomere of ChrR (ChrR:1,897,750 bp - 2,286,380 bp), reducing the dosage of 204 genes (**Fig 4E, S2A** and **Table S4**). GO analysis revealed that ChrRR-Deletion leads to a reduced dosage of 34/204 genes associated with the “response to stress” pathways and 18/204 genes linked to “response to drug” pathways (**Table S4**). We posit that this reduced gene dosage enables growth in the presence of high FLC. For example, *CKA1*, a gene whose deletion leads to FLC resistance [88], is located within the ChrRR-deletion (**Fig 4G**).

Interestingly, we found that all three *FLC-4* sequences colonies (*FLC-4a, b* and *c)*, are marked by a second segmental aneuploidy: a partial Chr1 amplification (Chr1-Duplication) (**Fig 4E** and **S2A**). This novel Chr1-Duplication amplifies a genomic fragment of ∼1.3 Mbp containing 535 protein-coding genes (**Table S4**). The Chr1-Duplication starts and ends near two distinct DNA repeat sequences with high sequence identity elsewhere in the genome: the 5’ breakpoint is within the *TLO34* and its 3’ breakpoint is within 3 kb of a *Zeta-1a* Long Terminal Repeat (LTR) (**Fig 4G** and **S3**) [32,33,89]. These WGS data led us to hypothesise that a chromosome-chromosome fusion event occurred between the Chr1-Duplication and Chr6 within homologous *TLO* sequences (**Fig 4G**). Indeed, the *TLO34* gene on Chr1 has high sequence identity with a 380 bp region located at Chr6 (position: 6182-6562 bp). In addition, sequence polymorphisms unique to Chr1-*TLO34* mapped to Chr6 in the *FLC-4* isolate (but not in *FLC-1, FLC-2* and *FLC-3*), supporting a novel interchromosomal recombination product between *TLO*-homologous sequences. This model is supported by CHEF gel electrophoresis analyses as, when compared to the *ulp2 Δ/Δ* progenitor, the *FLC-4* genome lacks one band corresponding to the shorter Chr6 homologue (blue asterisk), and it contains a new chromosome band of ∼2.2 Mb (magenta asterisk) (**Fig 4F**).

We posit that Chr1-Duplication provides a synergistic fitness advantage in response to two independent stressors (the presence of FLC and lack of *ULP2*) by simultaneously changing the dosage of several genes. Indeed, GO analyses demonstrated that 41 genes present in the Chr1-Duplication are associated with a” drug resistance” phenotypes (**Table S4**). Among these, amplification of *UPC2* encoding for the Upc2 transcription factor is likely to be critical. Indeed, it is well established that *UPC2* overexpression leads to FLC resistance by *ERG11* upregulation [90,91]. Chr1-Duplication likely rescues the fitness defects of the *ulp2 Δ/Δ* strain by amplifying two key genes: *CCR4* and *NOT5* (**Fig 4G**). Ccr4 and Not5 are subunits of the evolutionarily conserved Ccr4-Not complex that modulate gene expression at multiple levels, including transcription initiation, elongation, de-adenylation and mRNA degradation [92]. It has been shown that *S. cerevisiae CCR4* and *NOT5* overexpression rescue the lethal defects associated with a *ulp2* deletion strain [83].

Collectively our data suggest that the combined selective pressure of two independent stresses leads to selection of a chromosome aneuploidy that overcomes both stresses by overexpressing two different sets of genes.

## Discussion

In this study, we demonstrate that the SUMO protease Ulp2 is a critical regulator of *C. albicans* genome plasticity and that the development of drug resistance is accelerated in cells lacking *ULP2*. We unveil a striking flexibility of *C. albicans* cells in their response to complex stresses caused by drug treatment and dysregulation of the SUMO system, leading to the selection of extensive chromosome rearrangements.

### Ulp2 is a critical regulator of C. albicans genome stability

Our study identifies protein SUMOylation as a critical regulatory mechanism of *C. albicans* genome stability. SUMOylation is a dynamic and reversible post-translation modification in which a member of the SUMO family of proteins is conjugated to target proteins at lysine residues by E1 activating enzymes, E2 conjugating enzymes and E3 ligases [63–65]. SUMO is removed from its target proteins by SUMO-specific Ulp2 proteases [67]. Several observations are in agreement with our findings and suggest that SUMOylation controls stress-induced genome plasticity. Firstly, SUMOylation is a post-translational modification that is rapid and reversible, an essential requirement for a regulator of stress-induced genome plasticity. Secondly, *C. albicans* protein SUMOylation levels are different in normal and stress growth conditions [74]. Thirdly, deletion of genes encoding other components of the *C. albicans* SUMOylation machinery lead to filamentation, a phenotype often associated with defective cell division and compromised genome integrity [74,93,94]. Finally, *C. albicans* strains lacking the SUMO (Smt3) protein or the E3 ligase Mms21 display nuclear segregation defects [74,93].

*C. albicans* Ulp2 likely controls genome plasticity by modulating SUMO levels of several target proteins. SUMO proteases have a broad substrate specificity catalysing SUMO deconjugation of several substrates [95]. In other organisms, it is well known that SUMOylation modulates pathways ensuring genome integrity, including the DNA damage-sensing and repair pathway and the cell division and chromosome segregation pathway [63–66,96–98]. Despite the broad substrate specificity, our data suggest that one significant function of *C. albicans ULP2* is to ensure faithful chromosome segregation as high rates of chromosome missegregation is detected in the *ulp2 Δ/Δ* strain. Furthermore, the Illumina Genome sequencing analyses demonstrated that lack of *ULP2* is associated with extensive LOH events. Such extensive genomic changes are reminiscent of catastrophic mitotic events associated with defective chromosome segregation [99,100]. The targets of *C. albicans* Ulp2 are unknown, and it will be important to adopt proteomic approaches to identify the entire repertoire of SUMO targets and determine how *ULP2* contributes to *C. albicans* genome plasticity.

### Complex chromosome rearrangements drive adaptation to multiple stress environments

Our data demonstrate that the *ulp2 Δ/Δ* strain is more likely than the WT parental strain to develop resistance to anti-fungal drugs by selecting specific segmental aneuploidies on ChrR (ChrRR-deletion) and Chr1 (Chr1-duplication). These adaptive genotypes confer a growth advantage in response to two independent stressors: the absence of *ULP2* and drug treatment.

In agreement with the notion that repetitive elements play a significant role in genome instability, we identified intrachromosomal repetitive elements as drivers of genome instability. Indeed, all the sequenced *FLC*-adapted isolates carry a partial deletion of ChrR originating within the rDNA locus. We have previously demonstrated that the *C. albicans* rDNA locus is a hotspot for mitotic recombination [36], and clinical isolates are often marked by chromosomal aberrations originating from this locus [34]. This rDNA-driven chromosomal aberration leads to the deletion of one copy of 204 genes. We hypothesise that this reduced gene dosage drives FLC adaptation. For example, *CKA1*, one of the genes affected by ChrRR deletion, encodes for one of the two *C. albicans* Casein Kinases (Cka1 and Cka2). Deletion of these genes causes FLC resistance by controlling the expression of the efflux pump *CDR1* and *CDR2* [88].

WGS analysis demonstrated that the *FLC-4* isolate, whose FLC resistance is maintained followed by passaging on non-selective media, carries a second segmental aneuploidy: a partial duplication of Chr1 with breakpoints at repetitive elements. We provide evidence suggesting that Chr1 Duplication results from a fusion event between Chr1 and Chr6 due to a novel interchromosomal recombination product between *TLO* homologous sequences. We hypothesise that Chr1-duplication leads to gene dosage changes that are critical for overcoming two independent stresses: the presence of FLC and the absence of *ULP2*. Indeed, one of the master regulators of FLC resistance, *UPC2*, is located on the Chr1-duplication and its overexpression is likely to allow growth in the presence of FLC. *UPC2* encodes a key transcription factor of *ERG11*, the target of FLC [91]. It is well established that *UPC2* deletion leads to increased FLC susceptibility and that *UPC2* overexpression causes FLC resistance [91,101]. Accordingly, *UPC2* gain-of-function mutations are prevalent among FLC resistant clinical isolates [101].

The Chr1-duplication carries two key genes, *CCR4* and *NOT5*, likely to rescue the fitness defects associated with the *ulp2 Δ/Δ* strain. Indeed, it has been shown that *CCR4* and *NOT5* overexpression rescues the fitness defects of a *ULP2* deletion strain in *S. cerevisiae* [83]. Crr4 and Not5 are components of the evolutionarily conserved Crr4-Not multiprotein complex that regulate gene expression at all steps from transcription to translation and mRNA decay [102]. It is unknown why overexpression of the Crr4-Not complex rescues the fitness defect of an *ulp2* deletion strain, but it has been suggested that it might be linked to the transcriptional regulation of snoRNA and rRNA genes [84]. Here, for the first time, we demonstrate that segmental aneuploidy can lead to adaptation to different stressors by overexpressing genes located in the same chromosome and independently rescue the two stressors, leading to an overall fitness advantage.

## Material and Methods

### Yeast strains and Growth Conditions

Strains used in this study are listed in **Table S5**. Routine culturing was performed at 30 ºC in Yeast Extract-Peptone-D-Glucose (YPD) liquid and solid media containing 1% yeast extract, 2% peptone, 2% dextrose, 0.1 mg/ml adenine and 0.08 mg/ml uridine, Synthetic Complete (SC-Formedium) or Casitone (5 g/L Yeast extract, 9 g/L BactoTryptone, 20 g/L Glucose, 11.5 g/L Sodium Citrate dehydrate, 15 g/L Agar) media. When indicated, media were supplemented with 1mg/ml 5-Fluorotic acid (5-FOA, Melford), 200 μg/ml Nourseothricin (clonNAT, Melford), 5mM and 12 mM Caffeine (Sigma #C0750), 15 mg/ml and 128 mg/ml Fluconazole (Sigma #F8929), 6m H_2_O_2_ (Sigma #H1009), 12 mM and 22 mM Hydroxyurea (Sigma #H8627), 0.005% MMS (Sigma #129925).

### Genetic Screening

The genetic screening was performed using a *C. albicans* homozygous deletion library [44] arrayed in 96 colony format on YPD plates (145×20 mm) using a replica plater (Sigma #R2508). Control N/S plates were grown at 30 °C for 48 hours. UV treatment was performed using UVitec (Cambridge) with power density of 7.5µW/cm^2^ (0.030 J for 4 seconds). Following UV treatment, plates were incubated in the dark at 30°C for 48 hours. For MMS treatment, the library was spotted on YPD plates (145×20mm) containing 0.05% MMS and incubated at 30°C for 48 hours. UV and/or MMS sensitivity of selected strains was confirmed by serial dilution assays in control (YPD) and stress (UV: power density of 7.5µW/cm^2^, MMS: 0.05%) plates. Correct gene deletions were confirmed by PCR using gene-specific primers (**Table S6**).

### Yeast strain construction

Integration and deletion of genes were performed using long oligos-mediated PCR for gene deletion and tagging [103]. Oligonucleotides and plasmids used for strain constructions are listed in Supplementary **Table S6** and **S7**, respectively. For Lithium Acetate transformation, overnight liquid yeast cultures were diluted in fresh YPD and grown to OD_600_ of 1.3. Cells were harvested by centrifugation and washed once with dH_2_O and once with SORB solution (100mM Lithium acetate, 10mM Tris-HCL pH 7.5, 1mM EDTA pH 7.5/8, 1M sorbitol; pH 8). The pellet was resuspended in SORB solution containing single-stranded carrier DNA (Sigma-Aldrich) and stored -80 °C in 50 μl aliquots. Frozen competent cells were defrosted on ice, mixed with 5 µL of PCR product and 300 µL PEG solution (100mM Lithium acetate, 10mM Tris-HCL pH 7.5, 1mM EDTA pH 8, 40% PEG4000) and incubated for 21-24 hours at 30 °C. Cells were heat-shocked at 44°C for 15 minutes and grown in 5mL YPD liquid for 6 hours before plating on selective media at 30 °C.

### UV survival quantification

Following dilution of overnight liquid cultures, 500 cells were plated in YPD control plates while 1500 cells were plated in YPD stress plates and UV irradiates with power density of 7.5 µW/cm^2^ (0.030 J for 4 seconds). Plates were kept in the dark and incubated at 30°C for 48 hours. Colonies were counted using a colony counter (Stuart Scientific). Experiments were performed in 5 biological replicates, and violin plots graphs were generated using R Studio (http://www.r-project.org/).

### Growth curve

Overnight liquid cultures were diluted to 60 cells/µL in 100µL YPD and incubated at 30 °C in a 96 well plate (Cellstar®, #655180) with double orbital agitation of 400 rpm using a BMG Labtech SPECTROstar nanoplate reader for 48 hours. When indicated, YPD media was supplemented with MMS (0.05%) and HU (22 mM). Graphs show the average of 3 biological replicates and error bars show the standard deviation.

### Serial dilution assay

Overnight liquid cultures were diluted to an OD_600_ of 4, serially diluted 1:5 and spotted into agar plates with and without indicated additives using a replica plater (Replica plater for 96-well plates, Sigma Aldrich, #R2383). Images of the plates were then taken using Syngene GBox Chemi XX6 Gel imaging system. Experiments were performed in 3 biological replicates

### Protein extraction and Western blotting

Yeast extracts were prepared as described [104] using 1 × 10^8^ cells from overnight cultures grown to a final OD_600_ of 1.5–2. Protein extraction was performed in the presence of 2% SDS (Sigma) and 4 M acetic acid (Fisher) at 90°C. Proteins were separated in 2% SDS (Sigma), 40% acrylamide/bis (Biorad, 161-0148) gels and transfer into PVDF membrane (Biorad) by semi-dry transfer (Biorad, Trans Blot SD, semi-dry transfer cell). Western-blot antibody detection was used using antibodies from Roche Diagnostics Mannheim Germany (Anti-HA, mouse monoclonal primary antibody (12CA5 Roche, 5 mg/ml) at a dilution of 1:1000, and anti-mouse IgG-peroxidase (A4416 Sigma, 0.63 mg/ml) at a dilution of 1:5000, and Clarity™ ECL substrate (Bio-Rad).

### URA3^+^ marker loss quantification

Strains were first streaked on –Uri media to ensure the selection of cells carrying the *URA3*^*+*^ marker gene. Parallel liquid cultures. grown for 16 hours at 30°C in YPD, were plated on synthetic complete (SC) plates containing 1□mg/ml 5-FOA (5-fluorotic acid; Sigma) and on non-selective SC plates/. Colonies were counted after 2□days of growth at 30°C, the frequency of the *URA3*^*+*^ marker loss was calculated using the formula *F* = *m*/*M*, where *m* represents the median number of colonies obtained on 5-FOA medium corrected by the dilution factor used and the fraction of culture plated and *M* the average number of colonies obtained on YPD corrected by the dilution factor used and the fraction of culture plated [80]. Statistical differences between results from samples were calculated using the Kruskal-Wallis test and the Mann-Whitney U test for *post hoc* analysis. Statistical analysis was performed and violin plots were generated using R Studio (http://www.r-project.org/).

### Microscopy

30 ml of yeast cultures (OD_600_=1) grown in SC were centrifuged at 2000 rpm for 5 minute and washed once with dH_2_O. Cells were fixed in 10ml of 3.7% paraformaldehyde (Sigma #F8775) for 15 minutes, washed twice with 10ml of KPO_4_/Sorbitol (100 mM KPO_4_, 1.2 M Sorbitol) and resuspended in 250 μl PBS containing 10 μg of Dapi. Cells were then sonicated and resuspended in a 1% low melting point agarose (Sigma Aldrich) before mounting under a 22mm coverslip of 0,17um thickness. Samples were imaged on a Zeiss LSM 880 Airyscan with a 63x/1.4NA oil objective. Airyscan images were taken with a relative pinhole diameter of 0.2 AU (airy unit) for maximal resolution and reduced noise. GFP was imaged with a 488nm Argon laser and 495-550 nm bandpass excitation filter, RFP with a 546nm solid-state diode laser and a 570nm long pass excitation filter. The Dapi channel was imaged on a PMT with standard pinhole of 1AU and brightfield image were captured on the trans-PMT with the same excitation laser of 405nm., Dapi and brightfield images were taken with the same pixel size and bit depth (16bit) as the airyscan images. Images were of a 42.7×42.7um field of view and with a 33 nm pixel size resolution. z-stacks were taken containing cells of z interval of 500nm. Airyscan Veena filtering was performed with the inbuilt algorithms of Zeiss Zen Black 2.3. Fiji scripts were written to automatically create a maximum intensity projection with standardised intensity scaling for the fluorescence images and overlay them with the best focus image of the brightfield picture. Experiments were performed in 3 biological replicates and >100 cells/replicate were counted.

### Drug Selection

Strains were incubated overnight in casitone liquid media at 30°C with shaking. 10^4^ cells were plated in small (10cm) casitone plates or plates containing: (*i*) 128 µg/mL DMSO (Fluconazole Control), (*ii*) 128 µg/mL Fluconazole or (*iii*) 12 mM Caffeine. Plates were incubated at 30°C for 7 days. Colonies able to grow on Fluconazole- or Caffeine-containing plates were streaked in non-selective plates and tested by spotting assay in casitone+ DMSO plates, casitone+Fluconazole or casitone+Caffeine plates. Following incubation at 30°C, plates were imaged using Syngene GBox Chemi XX6 Gel imaging system. Experiments were performed in 3 biological replicates.

### Whole-genome sequence analysis

All genome sequencing data have been deposited in the Sequence Read Archive under BioProject PRJNA781758, Genomic DNA was isolated using a phenol-chloroform extraction as previously described [29]. Paired-end (2 × 151 bp) sequencing was carried out by the Microbial Genome Sequencing Center (MiGS) on the Illumina NextSeq 2000 platform. Adaptor sequences and low-quality reads were removed using Trimmomatic (v0.33 LEADING:3 Trailing:3 SLIDINGWINDOW:4:15 MINLEN:36 TOPHRED33) [105]. Trimmed reads were mapped to the *C. albicans* reference genome (A21-s02-m09-r08) from the *Candida* Genome Database (http://www.candidagenome.org/download/sequence/C_albicans_SC5314/Assembly21/archive/C_albicans_SC5314_version_A21-s02-m09-r08_chromosomes.fasta.gz). Reads were aligned to the reference using BWA-MEM (v0.7.17) with default parameters [106]. The BAM files, containing aligned reads, were sorted and PCR duplicates removed using Samtools (v1.10 samtools sort, samtools rmdup) [107]. Qualimap (v2.2.1) analysed the BAM files for mean coverage of the reference genome; coverages ranged from 73.7x to 89.3x coverage [108]. Variant detection was conducted using the Genome Analysis Toolkit (Mutect, v2.2-25) [109]. Variants were annotated using SnpEff (V4.3) [110] using the SC5314 reference genome fasta and gene feature file above. Parental variants were removed, and all remaining variants were verified visually using the Integrative Genomic Viewer (IGV, v2.8.2) [111].

### Read depth and breakpoint analysis

Whole-genome sequencing data were analysed for copy number and allele ratio changes as previously described [32,33]. Aneuploidies were visualised using the Yeast Mapping Analysis Pipeline (YMAP, v1.0) [112]. BAM files aligned to the SC5314 reference genome as described above were uploaded to YMAP and read depth was determined and plotted as a function of chromosome position. Read depth was corrected for both chromosome-end bias and GC-content. The GBrowse CNV track and GBrowse allele ratio track identified regions of interest for CNV and LOH breakpoints, and more precise breakpoints were determined visually using IGV. LOH breakpoints are reported as the first informative homozygous position in a region that is heterozygous in the parental genome. CNV breakpoints were identified as described previously [32,33].

### Contour-clamped homogeneous electric field (CHEF) electrophoresis

Intact yeast chromosomal DNA was prepared as previously described [113]. Briefly, cells were grown overnight, and a volume equivalent to an OD_600_ of 7 was washed in 50 mM EDTA and resuspended in 20 µl of 10 mg/ml Zymolyase 100T (Amsbio #120493-1) and 300 µl of 1% Low Melt agarose (Biorad® # 1613112) in 100 mM EDTA. Chromosomes were separated on a 1% Megabase agarose gel (Bio-Rad) in 0.5X TBE using a CHEF DRII apparatus. Run conditions as follows: 60-120s switch at 6 V/cm for 12 hours followed by a 120-300s switch at 4.5 V/cm for 12 hours, 14 °C. The gel was stained in 0.5x TBE with ethidium bromide (0.5 µg/ml) for 30 minutes and destained in water for 30 minutes. Chromosomes were visualised using a Syngene GBox Chemi XX6 gel imaging system.

## Acknowledgments

We thank Judith Berman for reagents, strains and materials and A. Pidoux for critical reading of the manuscript. This work was supported by BBSRC (BB/T006315/1 to A.B and S.V.E), a University of Kent GTA PhD studentships (to M.R.), a University of Minnesota UMR Fellowship with the Bioinformatics and Computational Biology program (to N.S), the National Institutes of Health (R01AI143689) and Burroughs Wellcome Fund Investigator in the Pathogenesis of Infectious Diseases Award (#1020388) to A.S.

## Notes

### Competing Interest Statement

The authors have declared no competing interest.

