## Supplementary material for "The SUMO protease Ulp2 regulates genome stability and drug resistance in the human fungal pathogen *Candida albicans*": Fig S1, Fig S2, Fig S3

**[Supporting Information](https://journals.plos.org/plosbiology/s/supporting-information): Figures**

*Rizzo et al*

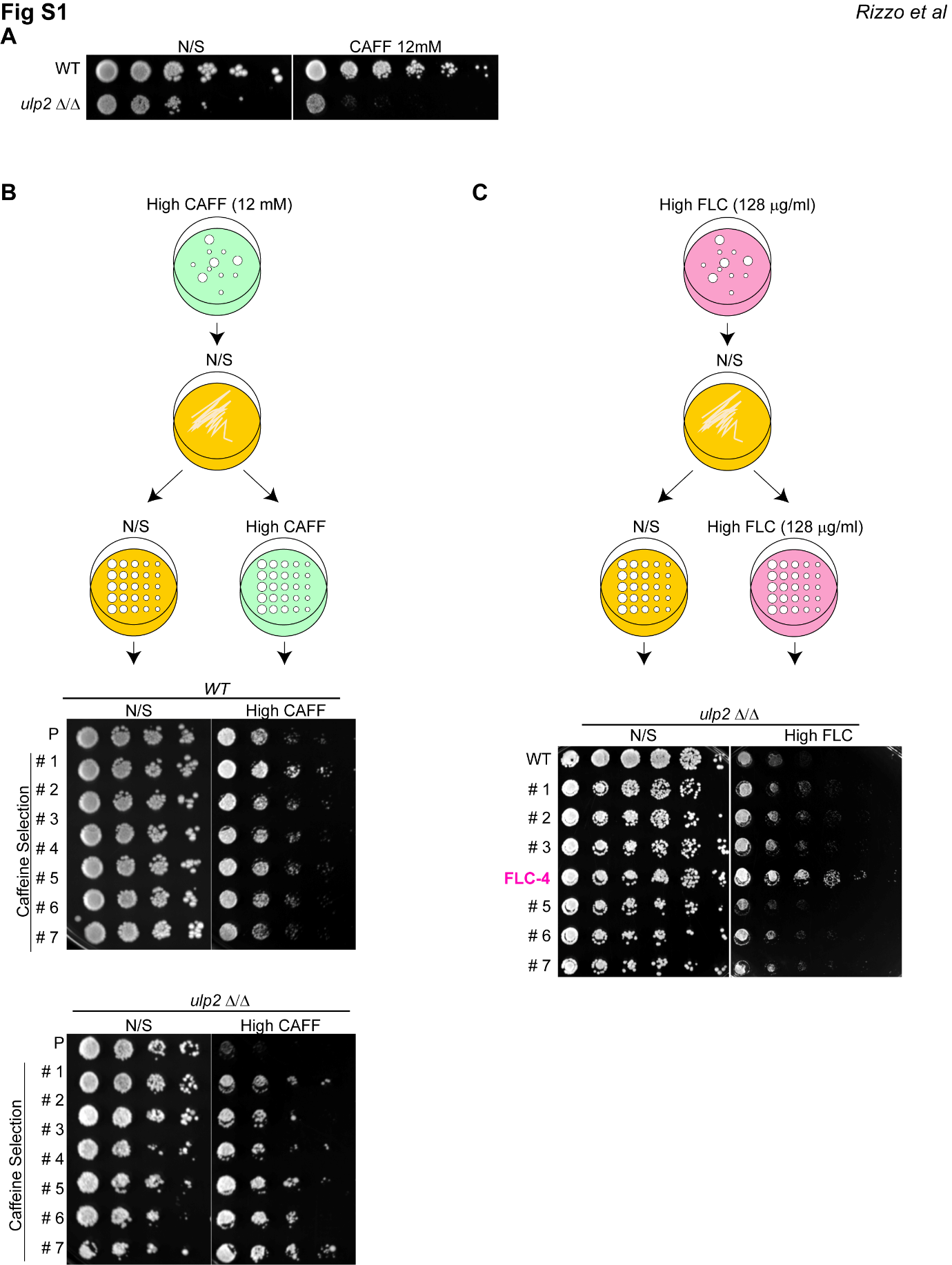
**Fig S1 (A**) Serial dilution assay of WT and *ulp2 Δ/Δ* strains grown in non-selective (N/S) media and media containing 12 mM Caffeine (CAF) **(B)** *Top:* Schematic of CAFF resistance testing. Single colonies from high CAF plates (12 mM) were streaked on non-selective (N/S) media before conducting a serial dilution assay in non/selective and High Caffein (12 mM) plates *Bottom:* Serial dilution assay of WT and *ulp2 Δ/Δ* colonies in non-selective media and High caffeine (12 mM CAF) media. Colonies were selected on high Caffeine (12 mM) and passaged in non-selective (N/S) media before performing the experiment. **(C)** *Top:* Schematic of FLC resistance testing. Single colonies from high FLC plates (128 μg/ml) were streaked on non-selective (N/S) media before conducting a serial dilution assay in non/selective and High FLC plates (128 μg/ml) *Bottom:* Serial dilution assay of WT and *ulp2 Δ/Δ* colonies in non-selective (N/S) media and High FLC (12 mM CAF) media. Colonies were selected on High FLC plates (128 μg/ml) and passaged in non-selective (N/S) media before performing the experiment. The sequenced *FLC-4* isolate is highlighted (magenta)

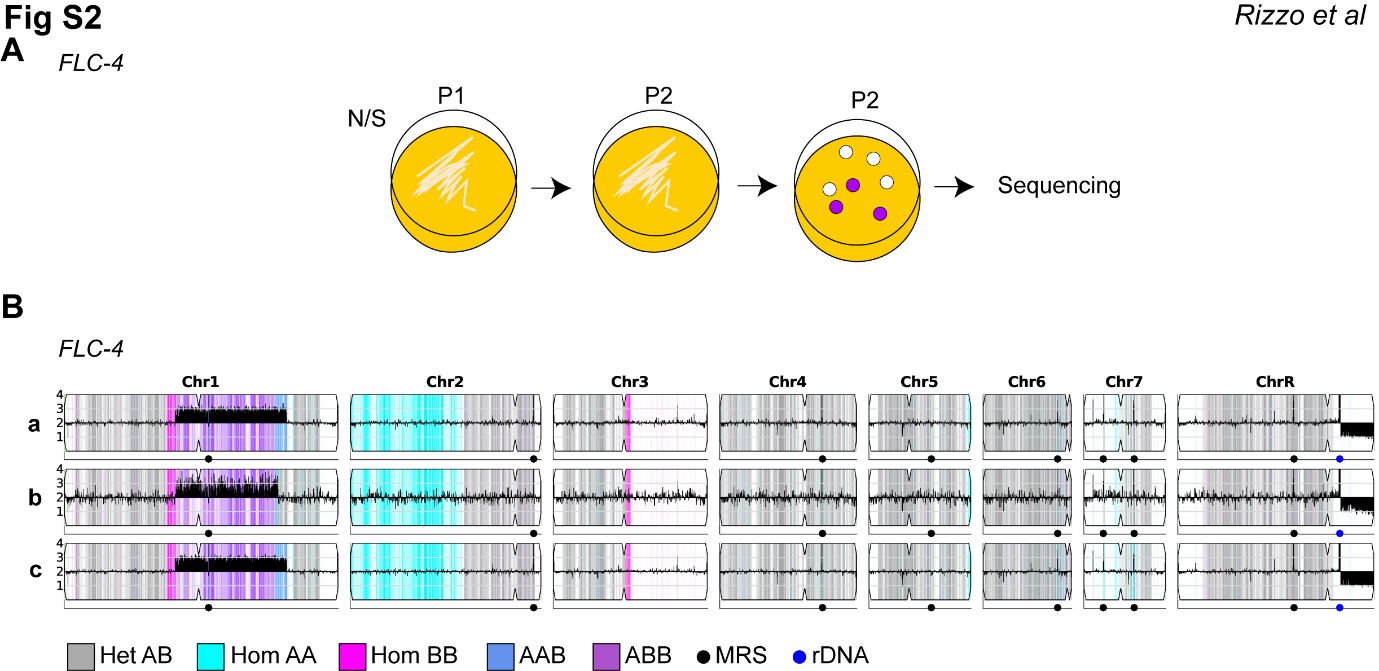

**Fig S2 (A)** Schematic of FLC-4 passaging before sequencing. 3 colonies (magenta) were sequenced. **(B)** FLC-4 was selected on 128 ug/ml FLC, passaged twice on YPAD, and then plated again for single colonies on 128 ug/ml FLC. Three single colonies (magenta) were selected and sent for whole genome sequencing. **(B)** Whole genome sequence data were plotted as the log2 ratio and converted to chromosome copy number (y-axis, 1-4 copies) as a function of chromosome position (x-axis, Chr1-ChrR) using YMAP. Heterozygous (AB) regions are indicated with gray shading and homozygous regions are indicated by haplotype AA (cyan) or BB (magenta). Allele ratio changes that occur within a CNV are indicated as dark blue (AAB) or purple (ABB). Colony B and C had allele ratio colouring that was corrected using IGV and allele frequency information. Two homozygous regions were already present in the progenitor (the left side of Chr2 and a small region near the centromere of Chr3).

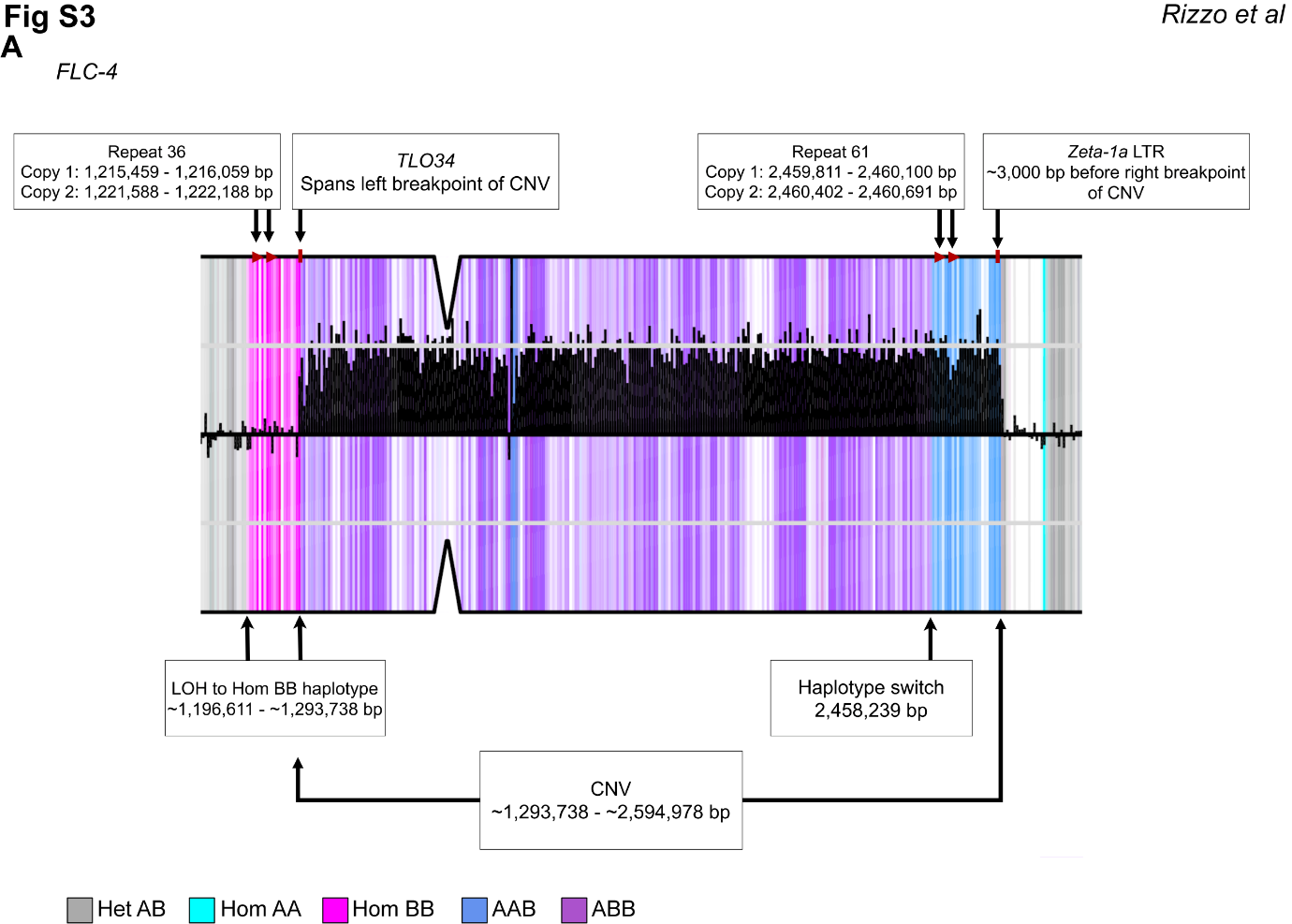
**Fig S3. Copy number and allele ratio changes occur near repeat sequences in FLC-4.** Whole genome sequence data plotted as in Figure S2 for the segmental amplification of Chr1 in one representative strain (FLC-4-c). Repetitive sequences are identified at the CNV and allele ratio changes. The position of copy number and allele ratio changes are approximate and repeat numbers refer to Supplementary file 2 from Todd et al., 2019. From left to right across Chr1: the alleles change from heterozygous (AB, gray) to homozygous (BB, magenta) at position 1,196,611 bp, near Repeat 36. The copy number increases from 2 to 3 copies, and the allele ratio changes to ABB (purple), at the repeat containing *TLO34*. On the other side of the CNV, there is a haplotype switch from purple (ABB) to dark blue (AAB) near repeat 61. Finally, the copy number decreases from 3 to 2 copies at the repeat containing Zeta-1a, at which point the allele ratio changes back to AB (gray).
