## Supplementary material for "The SUMO protease Ulp2 regulates genome stability and drug resistance in the human fungal pathogen *Candida albicans*": Table S5

**Table S5: Strains used in this study**

| **Strain number** | **Name** | **Genotype** | **Source** |
| --- | --- | --- | --- |
| AB55 | *SC5314* | WT | [1] |
| AB54 | *SN152* | *MTL a/α ura3∆-iro1∆::imm434/URA3^+^-IRO1 his1∆/his1∆ arg4∆/arg4∆ leu2∆/leu2∆* | [2] |
| AB140 | *TetR-GFP, TetO-CEN7* | *ORF19.1963::TetR-GFP-Nat::ORF19.1963/ORF19.1963 TetO-HIS::CEN7* | [3] |
| AB653 | *PHO85/pho85Δ::URA3^+^* | *ura3Δ::λimm434/ura3Δimm434 his1::hisG/his1::hisG PHO85/pho85* Δ*::URA3^+^* | [4] |
| AB655 | *CLB4/clb4∆::URA3^+^* | *ura3Δ::λimm434/ura3Δimm434 his1::hisG/his1::hisG CLB4/clb4 Δ::URA3* | [4] |
| AB663 | *CRZ1/CRZ1::GFP-URA3^+^* | *ura3Δ::λimm434/ura3Δimm434 his1::hisG/his1::hisG CRZ1/CRZ1::GFP-URA3* | [4] |
| AB755 | *ulp1∆/∆* | *MTL a/α ura3∆-iro1∆::imm434/URA3-IRO1 his1∆/his1∆ arg4∆/arg4∆ leu2∆/leu2∆ ulp1∆::NATR/ulp1∆::ARG^+^* | This work |
| AB758 | *ulp2∆/∆* | *MTL a/α ura3∆-iro1∆::imm434/URA3-IRO1 his1∆/his1∆ arg4∆/arg4∆ leu2∆/leu2∆ ulp2∆::NATR/ulp3∆::ARG^+^* | This work |
| AB746 | *SN250* | *his1Δ/his1Δ, leu2Δ::C.dub HIS1^+^ /leu2Δ::C.maltosa LEU2^+^, arg4Δ /arg4Δ, URA3/ura3Δ::imm^434^, IRO1/iro1Δ::imm^434^* | [2,5] |
| AB765 | *TetR-GFP, TetO-CEN7, ulp2∆/ulp2∆* | *ORF19.1963::TetR-GFP-Nat::ORF19.1963/ORF19.1963 TetO-HIS::CEN7 ulp2 ∆::ARG^+^ /ulp2∆::NATR* | This work |
| AB803 | *CLB4/clb4::URA3^+^ ulp2∆/ulp2∆* | *ura3Δ::λimm434/ura3Δimm434 his1::hisG/his1::hisG CLB4/clb4∆::URA3^+^ ulp2∆::NATR / ulp2∆::HIS^+^* | This work |
| AB804 | *CRZ1/CRZ1::GFP-URA3^+^ ulp2∆/ulp2∆* | *ura3Δ::λimm434/ura3Δimm434 his1::hisG/his1::hisG CRZ1/CRZ1::GFP-URA3^+^* *ulp2∆::NATR* / *ulp2∆::HIS^+^* | This work |
| AB823 | *PHO85/pho85::URA3^+^ ulp2∆/ulp2∆* | *ura3Δ::λimm434/ura3Δimm434 his1::hisG/his1::hisG PHO85/pho85∆:: URA3^+^ ulp2∆::NATR / ulp2∆::HIS^+^* | This work |
