## Supplementary material for "The SUMO protease Ulp2 regulates genome stability and drug resistance in the human fungal pathogen *Candida albicans*": Table S6

**Table S6: Oligonucleotides used in this study**

| **Sequence** | **Description** | **Name** | **AB Number** |
| --- | --- | --- | --- |
| CAAATCCATCAATGGATCAG | Check presence arg cassette | Arg4_Rev1 | 251 |
| CTGGTTGGAACAGAAGATTG | Check presence NAT cassette | Nat_Fw | 857 |
| ACTACCTGTTTCTGAACTAT | Check *MEC3* gene deletion | mec3_fw | 885 |
| CATCAGTATCATCTAGCAAG |  | mec3_rv | 886 |
| TGGAATCGAAGCAAGAGGTA | Check *RAD18* gene deletion | rad18_fw | 891 |
| ACCAAACGACGGTTGATGTT |  | rad 18 rv | 892 |
| ACTCATCACTGCGATATCAG | Check *ULP2* gene deletion | Ulp2_Fw | 905 |
| AGCTTCCATATGAGGAATCAA |  | Ulp2_Rv | 906 |
| ATGGTTCCGTCCTCATATGC | Check *GRR1* gene deletion | grr1_fw | 913 |
| ATGCATCTGTTATCTGCATACA |  | grr1_rv | 914 |
| GAAAAAAAAACAATACCAGCCATGATGAAGATTGCTTACACTATTTCTTCTTTATAACTGGACGTTACTAAgttttcccagtcacgacgtt | *ULP2* gene deletion with Arg cassette | Ulp2_Arg_Fw | 1032 |
| CATTTTTAGGTATCAAGTTTTGAAAAAAGAAAACAACAGTGGATGTGATATATATATAATCTTGTACATACAATTAGtggaattgtgagcggataa |  | Ulp2_arg_Rv | 1033 |
| ATTGGAAGAGGAATTGGAGA | Check *ULP2* gene deletion | Ulp2_check_Rv | 1034 |
| GGGGAAAAAAAAACAATACCAGCCATGATGAAGATTGCTTACACTATTTCTTCTTTATAACTGGACGTTACTAAgtaaaacgacggccagtgaa | *ULP2* gene deletion with NAT cassette | Ulp2_nat_Fw | 1038 |
| GGTATCAAGTTTTGAAAAAAGAAAACAACAGTGGATGTGATATATATATAATCTTGTACATACAATTAtgcatcaattgacgttgataccac |  | Ulp2_nat_Rv | 1039 |
| GTATAGTCAAAATGATATGAAAATAATTCGTAGAAGAATGGTCTATGAAATTTTAGATAATCGTTTACTAGATcggatccccgggttaattaa | HA tagging of *ULP2* gene | Ulp2_HA_Nat_Fw | 1042 |
| TAGGTATCAAGTTTTGAAAAAAGAAAACAACAGTGGATGTGATATATATATAATCTTGTACATACAATTAgtaaaacgacggccagtgaattc |  | Ulp2_HA_Nat_Rev | 1043 |
| CGTCCTTTCAAGTATTGTAACTGCCACGGACCAGACGAATGTTGAACTTTTGAAAATAAGATTAGAAAATgtaaaacgacggccagtgaat | *ULP1* gene deletion with NAT cassette | Ulp1_nat_fw | 1054 |
| TTACTATTGGATAAATACTACAGATAATTCTGATTGCTAGAATGTAGATATATATATAAATAAATTTAGTtgcatcaattgacgttgatac |  | Ulp1_nat_Rv | 1055 |
| CAAGATTGCAGATGCTTGAG | Check *ULP1* gene deletion | Ulp1_check_Rv | 1056 |
| GTCCTTTCAAGTATTGTAACTGCCACGGACCAGACGAATGTTGAACTTTTGAAAATAAGATTAGAAAATgttttcccagtcacgacgttgt | *ULP1* gene deletion with arg cassette | Ulp1_arg_Fw | 1057 |
| TTACTATTGGATAAATACTACAGATAATTCTGATTGCTAGAATGTAGATATATATATAAATAAATTTAGTgtggaattgtgagcggataac |  | Ulp1_arg_rv | 1058 |
