## Supplementary material for "The SUMO protease Ulp2 regulates genome stability and drug resistance in the human fungal pathogen *Candida albicans*": Table S7

**Table S7: Plasmids used in this study**

| **Plasmid** | **Description** | **AB Number** | **Source** |
| --- | --- | --- | --- |
| pHA_NAT | NAT substitution cassette  HA tagging | AB17 | [1] |
| pR3Arg46spe1 | ARG substitution cassette | AB18 | [2] |
| pGEM-His1 | HIS substitution cassette | AB20 | [2] |
